# The isoforms of pyruvate kinase act as nutrient sensors for the β-cell K_ATP_ channel

**DOI:** 10.1101/2022.02.09.478817

**Authors:** Hannah R. Foster, Thuong Ho, Evgeniy Potapenko, Sophia M. Sdao, Sophie L. Lewandowski, Halena R. VanDeusen, Shawn M. Davidson, Rebecca L. Cardone, Marc Prentki, Richard G. Kibbey, Matthew J. Merrins

**Affiliations:** Department of Medicine, Division of Endocrinology, Diabetes, and Metabolism, University of Wisconsin-Madison, Madison, WI, USA; Koch Institute for Integrative Cancer Research, Massachusetts Institute of Technology, Cambridge, MA, USA; Lewis-Sigler Institute for Integrative Genomics, Princeton University, Princeton, NJ, USA; Department of Internal Medicine, Yale University, New Haven, CT, USA; Molecular Nutrition Unit and Montreal Diabetes Research Center, CRCHUM, and Departments of Nutrition, Biochemistry and Molecular Medicine, Université de Montréal, Montréal, Canada; Department of Cellular & Molecular Physiology, Yale University, New Haven, CT, USA; William S. Middleton Memorial Veterans Hospital, Madison, WI, USA

**Keywords:** Pyruvate kinase, phosphoenolpyruvate, PKm1, PKm2, PCK2, K_ATP_ channels, β-cell metabolism, PEP cycle, ATP/ADP, Ca^2+^ oscillations, insulin secretion

## Abstract

Pyruvate kinase (PK) and the phosphoenolpyruvate (PEP) cycle play key roles in nutrient-stimulated K_ATP_ channel closure and insulin secretion. To identify the PK isoforms involved, we generated mice lacking β-cell PKm1, PKm2, and mitochondrial PEP carboxykinase (PCK2) that generates mitochondrial PEP. Glucose metabolism generates both glycolytic and mitochondrially-derived PEP, which triggers K_ATP_ closure through local PKm1 and PKm2 signaling at the plasma membrane. Amino acids, which generate mitochondrial PEP without producing glycolytic fructose 1,6-bisphosphate to allosterically activate PKm2, signal through PKm1 to raise ATP/ADP, close K_ATP_ channels, and stimulate insulin secretion. Raising cytosolic ATP/ADP with amino acids is insufficient to close K_ATP_ channels in the absence of PK activity or PCK2, indicating that K_ATP_ channels are regulated by mitochondrially-derived PEP that provides ATP via plasma membrane-associated PK, but not via mitochondrially-derived ATP. Following membrane depolarization, the PEP cycle is also involved in an “off-switch” that facilitates K_ATP_ channel reopening and Ca^2+^ extrusion, as shown by PK activation experiments and β-cell PCK2 deletion that prolonged Ca^2+^ oscillations and increased insulin secretion. In conclusion, the differential response of PKm1 and PKm2 to the glycolytic and mitochondrial sources of PEP influences the β-cell nutrient response, and controls the oscillatory cycle regulating insulin secretion.

## INTRODUCTION

Maintenance of euglycemia relies on β-cells to couple nutrient sensing with appropriate insulin secretion. Insulin release is stimulated by the metabolism-dependent closure of ATP-sensitive K^+^ (K_ATP_) channels (Ashcroft et al., 1984; Cook and Hales, 1984; Misler et al., 1986; Rorsman and Trube, 1985), which triggers Ca^2+^ influx and exocytosis (Anderson and Long, 1947; Grodsky et al., 1963). Contrary to what is often believed, the glucose-induced signaling process in β-cells has not been largely solved, and the entrenched model implicating a rise in mitochondrially-derived ATP driving K_ATP_ channel closure (Campbell and Newgard, 2021; Prentki et al., 2013) is incomplete and possibly wrong, the main reason being that it does not consider other sources of local ATP production that may be key for signaling (Corkey, 2020; Lewandowski et al., 2020). The recent discovery that pyruvate kinase (PK), which converts ADP and phosphoenolpyruvate (PEP) to ATP and pyruvate, is present on the β-cell plasma membrane where it is sufficient to raise sub-plasma membrane ATP/ADP (ATP/ADP_pm_) and close K_ATP_ channels (Lewandowski et al., 2020) provides an alternative mechanism to oxidative phosphorylation for K_ATP_ channel regulation. Based on this finding, Lewandowski *et al*. proposed a revised model of β-cell fuel sensing, which we refer to here as the Mito_Cat_-Mito_Ox_ model, that is relevant to both rodent and human islets and the *in vivo* context (Abulizi et al., 2020; Lewandowski et al., 2020).

In the Mito_Cat_-Mito_Ox_ model of β-cell metabolic signaling, Ca^2+^ and ADP availability dictate the metabolic cycles that preferentially occur during the triggering or secretory phases of glucose-stimulated oscillations (Figure S1). The triggering phase, referred to as Mito_Cat_ (a.k.a. Mito_Synth_ (Lewandowski et al., 2020)), is named for the matched processes of anaplerosis (i.e., the net filling of TCA cycle intermediates) and cataplerosis (i.e., the egress of TCA cycle intermediates to the cytosol). During this electrically-silent phase of metabolism, the favorable bioenergetics of PEP metabolism (ΔG° = −14.8 kcal/mol for PEP vs. −7.3 for ATP) by PK progressively increases the ATP/ADP ratio, and by lowering ADP slows oxidative phosphorylation. The shift to a higher mitochondrial membrane potential (ΔΨ_m_) elevates the NADH/NAD^+^ ratio in the mitochondrial matrix and slows the TCA cycle, increasing acetyl-CoA that allosterically activates pyruvate carboxylase, the anaplerotic consumer of pyruvate that fuels oxaloacetate-dependent PEP synthesis by mitochondrial PEP carboxykinase (PCK2). The return of mitochondrial PEP to the cytosol completes the “PEP cycle” that helps fuel PK, which raises ATP/ADP_pm_ to close K_ATP_ channels. Following membrane depolarization and Ca^2+^ influx, the increased workload (ATP hydrolysis) associated with ion pumping and exocytosis elevates cytosolic ADP, which activates oxidative phosphorylation to produce ATP that sustains insulin secretion in a phase referred to as Mito_Ox_. An unresolved aspect of this model is whether plasma membrane-compartmentalized PK activity is strictly required to close K_ATP_ channels. This question is important because in the current canonical model of fuel-induced insulin secretion, an increase in the bulk cytosolic ATP/ADP ratio (ATP/ADP_c_) is generally assumed to close K_ATP_ channels.

In the Mito_Cat_-Mito_Ox_ model, PK has two possible sources of PEP that may differentially regulate K_ATP_ closure: glycolytic PEP produced by enolase, and mitochondrial PEP produced by PCK2 in response to anaplerosis (Figure 1a). About 40% of glucose-derived PEP is generated by PCK2 in the PEP cycle and is closely linked to insulin secretion (Abulizi et al., 2020; Jesinkey et al., 2019; Stark et al., 2009). However, it remains unclear how the PEP cycle influences glucose-stimulated oscillations. Mitochondrial PEP derived from PCK2 may provide a glycolysis-independent mechanism by which PK rapidly increases ATP/ADP_pm_ locally at the K_ATP_ channel in response to amino acids, which are potent anaplerotic fuels.

**Figure 1.**
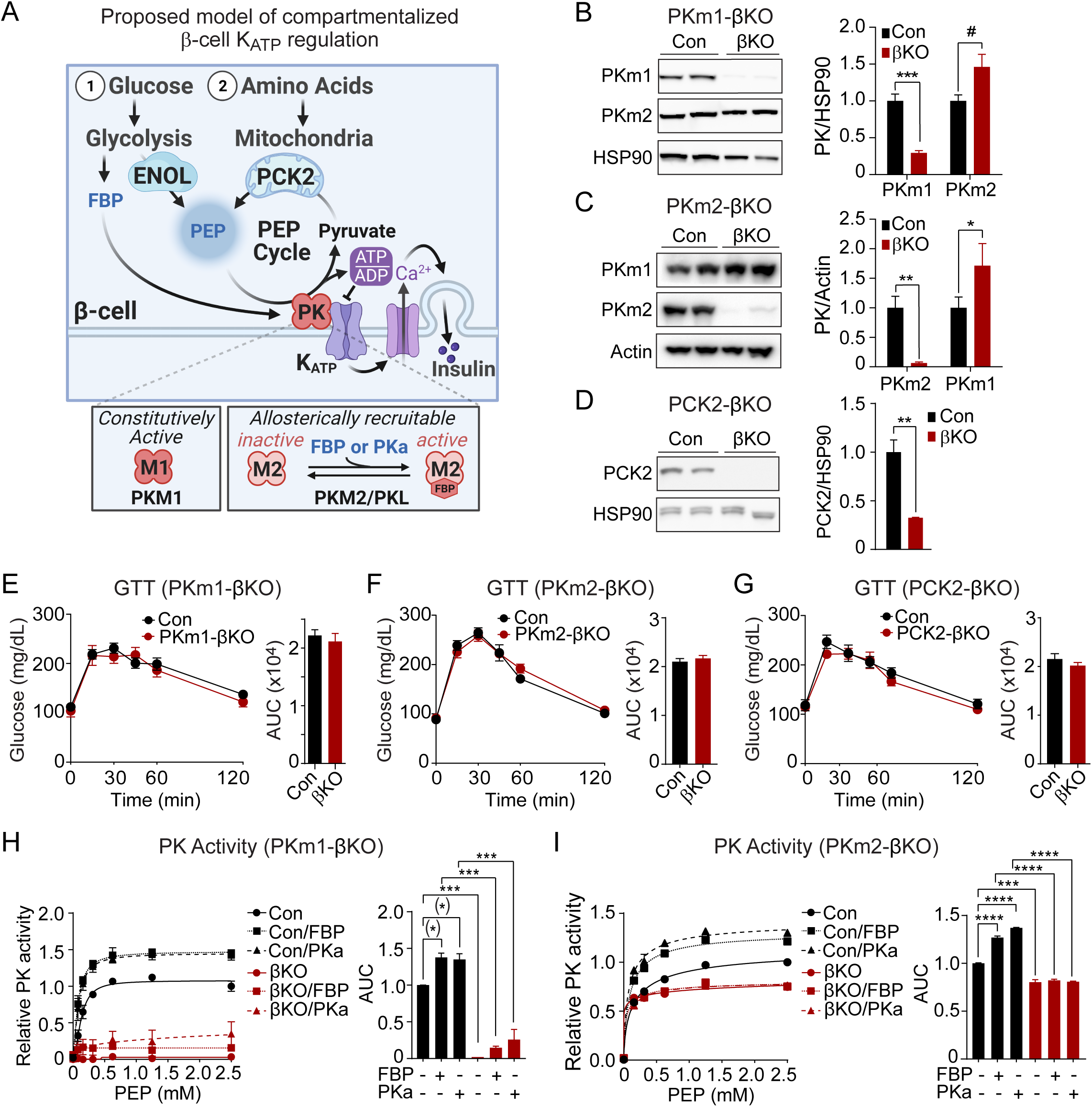
Generation of mouse models to probe the functions of PKm1, PKm2, and PCK2 in β-cells. Hypothesized model in which PK in the K_ATP_ microcompartment is fueled by two sources of PEP—glycolytic PEP generated by enolase, and mitochondrial PEP generated by PCK2 in response to anaplerotic fuels. β-cells express three isoforms of PK, constitutively active PKm1 and allosterically recruitable PKm2 and PKL that are activated by endogenous fructose 1,6-bisphosphate (FBP) or pharmacologic PK activators (PKa). (B-D) Quantification of knockdown efficiency in islet lysates from PKm1-βKO (B), PKm2-βKO (C), and Pck2-βKO mice (D) (*n* = 4 mice for PKm1- and PCK2-βKO, *n* = 6 mice for PKm2-βKO). (E-G) Intraperitoneal glucose tolerance tests (GTT, 1 g/kg) of PKm1-βKO mice (*n* = 9) and littermate controls (*n* = 8) (E), PKm2-βKO mice (*n* = 7) and littermate controls (*n* = 7) (F), and Pck2-βKO mice (*n* = 10) and littermate controls (*n* = 7) (G) following an overnight fast. (H-I) PK activity in islet lysates of PKm1-βKO (H) and PKm2-βKO mice (I) in response to FBP (80 μM) and PKa (10 μM TEPP-46). (*n* = 2 replicates from 6 mice/group) Data are shown as mean ± SEM. ^#^*P* < 0.01, \**P* < 0.05, \*\**P* < 0.01, \*\*\**P* < 0.001, **** *P* < 0.0001 by t-test (B-G) or two-way ANOVA (H-I).

The isoforms of PK, each with different activities and mechanisms of control, may differentially regulate K_ATP_ channels (Figure 1a). β-cells express the constitutively-active PKm1 as well as two allosterically-recruitable isoforms, PKm2 and PKL, which are activated by glycolytic fructose-1,6-bisphosphate (FBP) generated upstream by the phosphofructokinase reaction (DiGruccio et al., 2016; MacDonald and Chang, 1985; Mitok et al., 2018). Pharmacologic PK activators (PKa), which lower the K_m_ of PKm2 for PEP and increase the V_max_ (Anastasiou et al., 2012), increase the frequency of glucose-stimulated Ca^2+^ and ATP/ADP oscillations and potentiate nutrient-stimulated insulin secretion from rodent and human islets (Abulizi et al., 2020; Lewandowski et al., 2020). Much less is known about the PKm1 isoform, which due to its constitutive (FBP-insensitive) activity might be ideal in situations of high oxidative workload, as in cardiac myocytes (Li et al., 2021). β-cells may shift their reliance upon different PK isoforms throughout the oscillatory cycle, as the levels of glycolytic FBP rise during Mito_Cat_ and fall during Mito_Ox_ (Lewandowski et al., 2020; Merrins et al., 2016, 2013).

Here, we show that PK is essential for K_ATP_ closure – amino acids that effectively raise ATP/ADP_c_ cannot close K_ATP_ channels without PK. We further demonstrate that both PKm1 and PKm2 are active in the K_ATP_ channel microcompartment with at least two required functions. First, spatial privilege provides redundancy in the β-cell glucose response, by permitting the minor PKm2 isoform, when activated by FBP, to transmit the signal from glucose to K_ATP_ despite contributing only a small fraction of the whole cell PK activity. Second, the composition of PK isoforms within the K_ATP_ compartment tunes the β-cell response to amino acids, which provide mitochondrial PEP for PKm1 without also generating the FBP needed to allosterically activate PKm2. Using β-cell PCK2 deletion, we found that mitochondrially-derived PEP signals to the plasma membrane PK-K_ATP_ microcompartment during Mito_Cat_, and facilitates Ca^2+^ extrusion during Mito_Ox_. These studies support the Mito_Cat_-Mito_Ox_ model of oscillatory metabolism, and identify unique functions of the PKm1- and PKm2-driven PEP cycles in β-cell nutrient signaling.

## RESULTS

### PKm1 accounts for >90% of total β-cell PK activity, with <10% from PKm2 and no discernable contribution from PKL

We generated β-cell specific PKm1 and PKm2 knockout mice by breeding *Ins1-Cre* mice (Thorens et al., 2015) with *Pkm1*^*f/f*^ mice (Davidson et al., 2021; Li et al., 2021) (PKm1-βKO) or *Pkm2*^*f/f*^ mice (Israelsen et al., 2013) (PKm2-βKO). PKm1 protein was reduced by 71% in PKm1-βKO islets, while PKm2 increased by 46% compared to littermate *Ins1-Cre* controls (Figure 1b). Expression of PKm2 protein fell by 94% in PKm2-βKO islets, while PKm1 increased by 29% (Figure 1c). In both strains this partial compensation is expected since PKm1 and PKm2 are alternative splice variants of the *Pkm* gene (Li et al., 2021; Israelsen et al., 2013). We generated *Pck2*^*f/f*^ mice (Figure S2) and crossed them with *Ucn3-Cre* mice to facilitate postnatal β-cell deletion without the need for tamoxifen (Adams et al., 2021; van der Meulen et al., 2017). Islet PCK2 protein dropped by 67% in the Pck2-βKO compared to Pck2-floxed littermate controls (Figure 1d). None of these knockout mice were glucose intolerant (Figure 1e-g).

The relative contributions of each PK isoform relative to total PK activity was determined in the islet lysates. The endogenous allosteric metabolite, FBP, and pharmacologic PK activators such as TEPP-46 (PKa), have no impact on PKm1 but substantially lower the K_m_ and raise V_max_ of PKm2 or PKL (Lewandowski et al., 2020). Control islet lysates had a K_m_ for PEP of ∼140 ± 14 μM that was reduced in the presence of exogenous FBP (K_m,_ 100 μM ± 10 μM) or PKa (K_m_, 90 ± 13 μM), while V_max_ increased by about one-third (control, 1.05 ± 0.019 μmol/min; FBP, 1.28 ± 0.006 μmol/min; PKa, 1.36 ± 0.006 μmol/min). Quite remarkably, the V_max_ for islet PK activity from PKm1-βKO mice decreased by 97% compared to controls (Figure 1h), and was too low to estimate K_m_ accurately. The residual PK remained sensitive to activation by both FBP and PKa, identifying an allosterically-recruitable PK pool that accounts for only about 10% of the PK activity present in control islets (Figure 1h), despite the fact that PKm2 protein is elevated by 46% in the PKm1-βKO (Figure 1b). Conversely, β-cell PKm2 deletion lowered islet lysate PK V_max_ by ∼20% in the absence of activators, and eliminated both the K_m_ and V_max_ response to PKa and FBP (Figure 1i), thus ruling out any significant PKL activity. Taken together, mouse islet PK activity is composed of >90% PKm1, with a variable contribution from PKm2 depending on the FBP level. If only considered in terms of total cellular activity related to nutrient-induced insulin secretion (*i*.*e*. in the absence of any compartmentalized functions), PKm1 should be dominant over PKm2 under all physiologic conditions. The fact that PKm1-βKO mice maintain metabolic health with unaltered glucose tolerance into adulthood suggests that the remaining PK activity is sufficient for β-cell function, and led us to hypothesize that both PKm1 and PKm2 function in the K_ATP_ channel microcompartment.

### Both PKm1 and PKm2 are associated with the plasma membrane and locally direct K_ATP_ channel closure, however PKm2 requires allosteric activation

We previously demonstrated that PEP, in the presence of saturating ADP concentrations, can close K_ATP_ channels in mouse and human β-cells (Lewandowski et al., 2020). This suggests that PK is present near K_ATP_ and locally lowers ADP and raises ATP to close K_ATP_ channels, and that bulk ADP is of less importance than PEP for channel regulation. Excised patch-clamp experiments, which expose the inside of the plasma membrane to the bath solution (*i*.*e*. the inside-out mode), provide both the location of PK as well as its functional coupling to K_ATP_ channels in native β-cell membranes. This approach was applied in combination with β-cell deletion of PKm1 and PKm2 to directly identify the isoforms of the enzyme present in the K_ATP_ microdomain. K_ATP_ channels were identified by inhibition with 1 mM ATP, which blocked the spontaneous opening that occurs after patch excision (Figure 2a). Channel activity was restored using a test solution containing 0.5 mM ADP and 0.1 mM ATP. In control β-cells, the further addition of 5 mM PEP closed K_ATP_, as shown by a 77% reduction in the total power (a term reflecting both the frequency and channel open time) (Figure 2a), compared with only a 29% reduction in PKm1-βKO cells (Figure 2b). Note that K_ATP_ channel closure occurred in control β-cells despite the continuous deluge of the channel-opener ADP. Thus, it is not the PEP itself, but the PK activity in the K_ATP_ microcompartment that is responsible for K_ATP_ closure.

**Figure 2.**
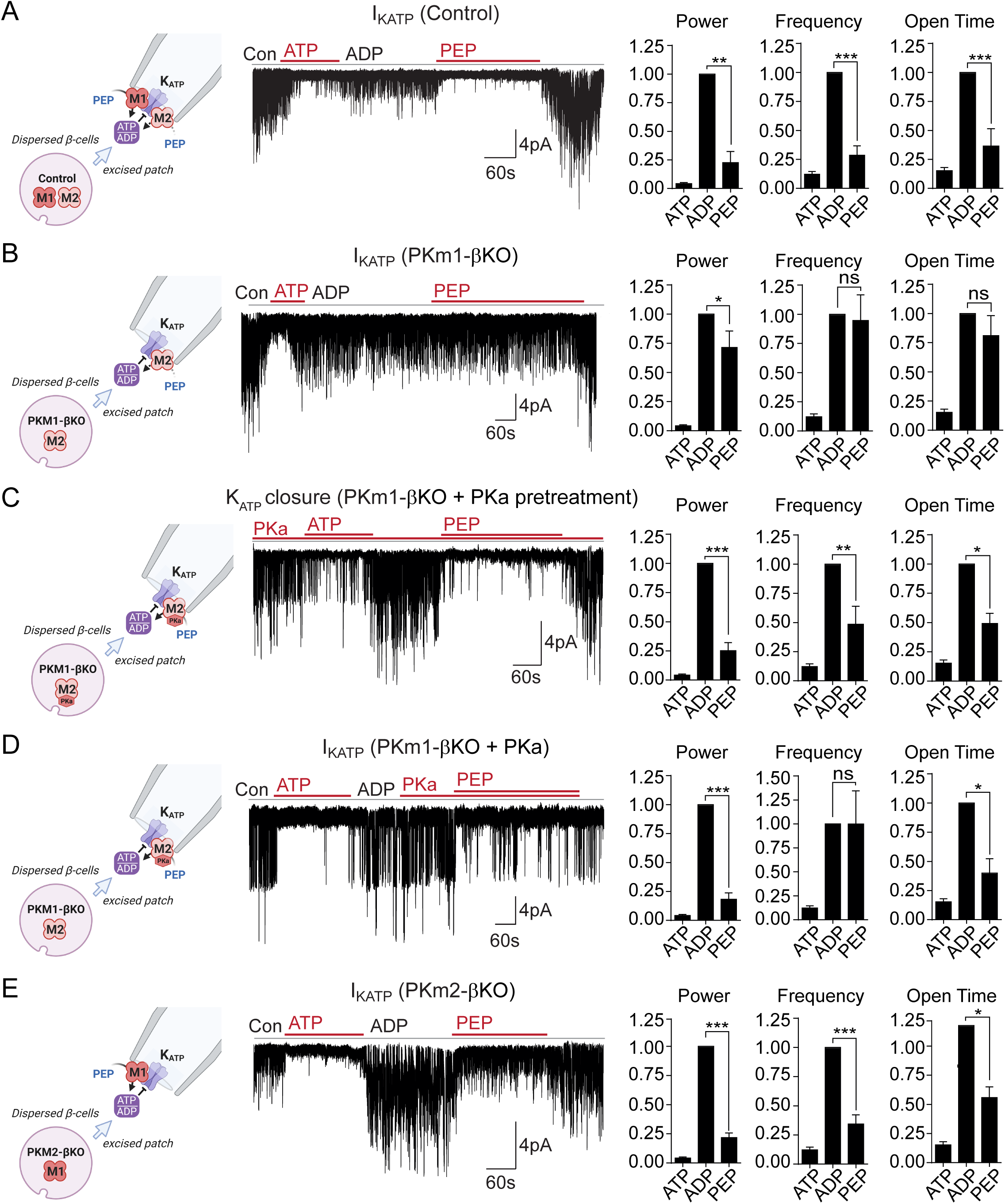
Plasma membrane K_ATP_ channels are locally regulated by a combination of PKm1 and allosterically activated PKm2. (A-E) K_ATP_ channel activity (holding potential = −50 mV) quantified in terms of power, frequency, and open time. Applying the substrates for PK closes K_ATP_ channels in excised patches of β-cell plasma membrane from control mice (*n* = 14 recordings from 4 mice) (A). Defective K_ATP_ channel closure in β-cells from PKm1-KO (*n* = 20 recordings from 5 mice) (B) is rescued by PKa pretreatment (*n* = 6 recordings from 3 mice) (C) and acute PKa application (*n* = 7 recordings from 3 mice) (D). PKm1 is sufficient for K_ATP_ closure in β-cells from PKm2-βKO (*n* = 6 recordings from 3 mice) (E). ATP, 1 mM; ADP, 0.5 mM ADP + 0.1 mM ATP; PEP, 5 mM; PKa, 10 μM TEPP-46. Data are shown as mean ± SEM. \**P* < 0.05, \*\**P* < 0.01, \*\*\**P* < 0.001, **** *P* < 0.0001 by paired t-test. ^(*)^*P* < 0.05 by unpaired t-test in (H).

To test for a role of PKm2 in the K_ATP_ microcompartment, PKm1-βKO cells were preincubated in the presence of 10 μM PKa, which restored PEP-dependent K_ATP_ channel closure to the same extent as the control (1 mM ATP) (Figure 2c). PKa had a similar effect when applied acutely (Figure 2d), indicating that PKm2 does not require allosteric activation to localize to the plasma membrane. Notably, the PEP concentration in the incubations were well over the K_m_ of either isoform. Therefore, the response of PKm2 to PKa at high PEP levels indicates that K_ATP_ closure requires an additional increase in the V_max_ via either allosteric activation of individual subunits of PKm2, or perhaps more likely, that the functional interaction is sensitive to the quaternary structure of PKm2. β-cells lacking PKm2 maintained channel closure with the same power as 1 mM ATP, which is attributable to the sufficiency of endogenous PKm1 (Figure 2e). Thus, metabolic compartmentation of PKm1 and PKm2 to the plasma membrane provides a redundant mechanism of K_ATP_ channel regulation, as well as a compelling explanation for the ability of PKm1-βKO mice to tolerate a near-complete loss of β-cell PK activity (Figure 1e,h).

### PKm1 and PKm2 are redundant for glucose-dependent Ca^2+^ influx

The rescue of PKm1 deficiency by PKa in the K_ATP_ microcompartment (Figure 2c-d) suggests that PKm1 and PKm2 exert redundant control over K_ATP_ closure, provided that glucose is present to generate FBP to activate PKm2. To test this further we examined Ca^2+^ dynamics with FuraRed while using a near-infrared dye, DiR, to facilitate simultaneous imaging of PKm1-, PKm2-, and PCK2-βKO islets with their littermate controls (Figure 3a). β-cell deletion of PKm1 or PKm2 did not reveal any biologically meaningful differences in the oscillatory period, the fraction of each oscillation spent in the electrically-active state (*i*.*e*. the duty cycle), or the amplitude of glucose-stimulated Ca^2+^ oscillations in intact islets (Figure 3b,d). In addition, PKm1 and PKm2 knockouts had no discernable difference in first-phase Ca^2+^ parameters (i.e., time to depolarization, amplitude, and duration of first phase) following an acute rise in glucose from 2 to 10 mM (Figure 3c,e). These data confirm *in situ* that PKm1 and PKm2 are redundant at high glucose.

**Figure 3.**
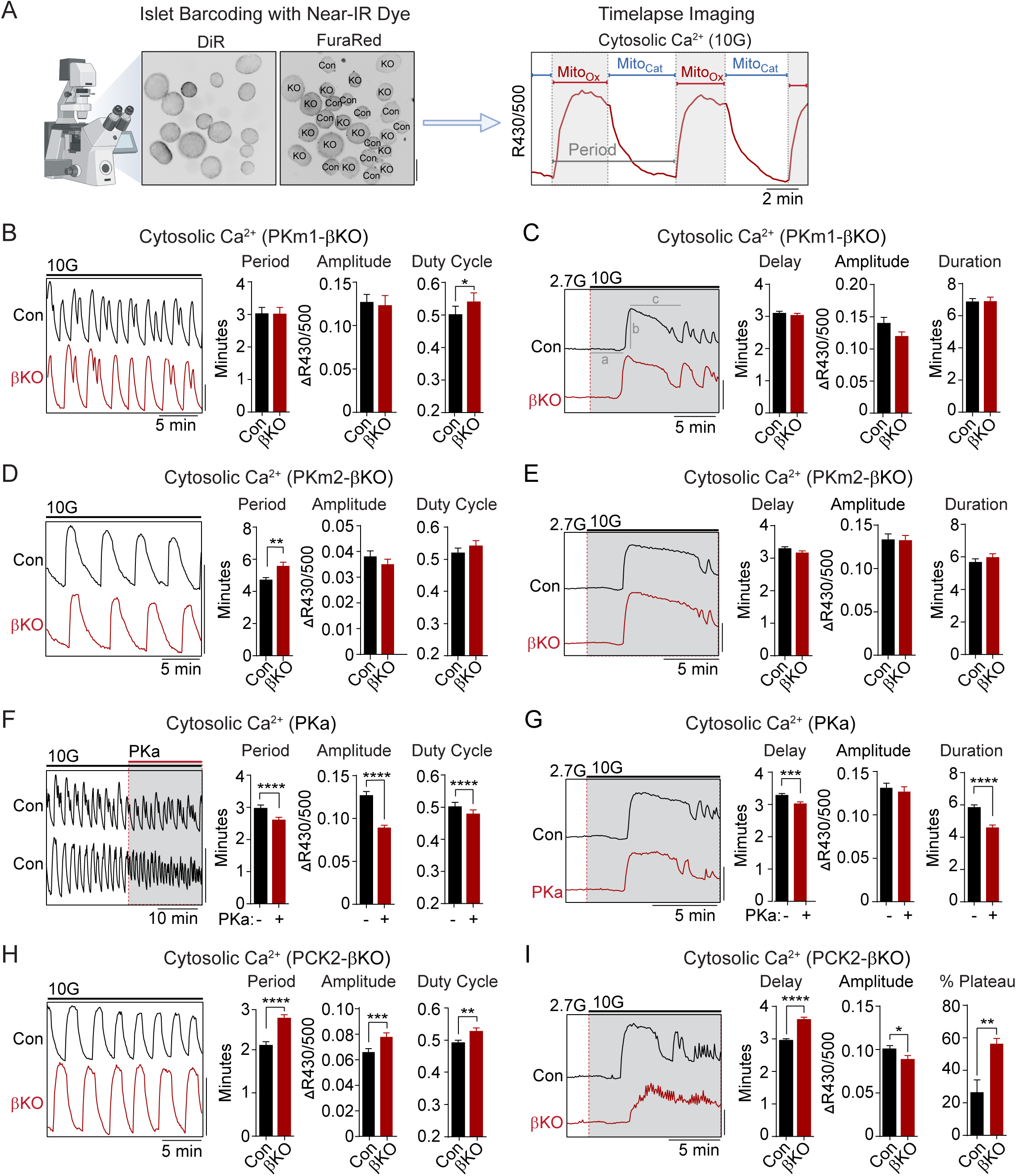
PKm2 and PCK2, but not PKm1, have metabolic control over first-phase and steady-state Ca_2+_ influx in response to glucose. (A) Barcoding of islet preparations with near-IR dye (DiR) permits simultaneous timelapse imaging of islet Ca^2+^ dynamics of control and βKO mice (scale bar = 200 μm) (*left*). A representative trace illustrates the cataplerotic triggering phase (Mito_Cat_) and oxidative secretory phases (Mito_Ox_) of steady-state Ca^2+^ oscillations in the presence of 10 mM glucose and 1 mM leucine (*right*). Gray line denotes the period. (B, D, F, H) Representative traces and quantification of period, amplitude, and duty cycle of steady-state Ca^2+^ oscillations in islets from PKm1-βKO (*n* = 91 islets from 3 mice) and littermate controls (*n* = 94 islets from 3 mice) (B), PKm2-βKO (*n* = 118 islets from 4 mice) and littermate controls (*n* = 111 islets from 4 mice) (D), control mice treated with PKa (10 μM TEPP-46) (*n* = 88 islets from 3 mice) (F), and PCK2-βKO (*n* = 74 islets from 3 mice) and littermate controls (*n* = 77 islets from 3 mice) (H). The bath solution (PSS) contained 10 mM glucose (10G) and 1 mM leucine. Scale bars: 0.1 FuraRed excitation ratio (R430/500). (C, E, G, I) Representative Ca^2+^ traces and quantification of time to depolarization (a), first-phase amplitude (b), and first-phase duration (c) in islets from PKm1-βKO (*n* = 144 islets from 6 mice) and littermate controls (*n* = 150 islets from 6 mice) (C), PKm2-βKO (*n* = 52 islets from 2 mice) and littermate controls (*n* = 55 islets from 2 mice) (E), PKa-treated (10 μM TEPP-46) (*n* = 161 islets from 9 mice) and vehicle controls (*n* = 212 islets from 9 mice) (G), and PCK2-βKO (*n* = 73 islets from 3 mice) and littermate controls (*n* = 78 islets from 3 mice) (I). The bath solution (PSS) contained 1 mM leucine and 2.7 mM (2.7G) and 10 mM glucose (10G) as indicated. Scale bars: 0.1 FuraRed excitation ratio (R430/500). Data are shown as mean ± SEM. \**P* < 0.05, \*\**P* < 0.01, \*\*\**P* < 0.001, **** *P* < 0.0001 by unpaired t-test (A-C and E-I) and paired t-test (D).

### PK and the PEP cycle are implicated in both on- and off-switches for Ca^2+^ influx

To study the interaction of PK with the PEP cycle, we performed islets Ca^2+^ measurements using PK activators and PCK2-βKO islets, in the latter case using islet barcoding to simultaneously image islets isolated from littermate controls. Consistent with the ability of allosteric PKm2 activation to accelerate K_ATP_ closure, acute application of PKa to wild-type islets reduced the period as well as the amplitude of the steady-state glucose-induced Ca^2+^ oscillations (Figure 3f and Figure S3a). However, we noticed that PKa shortened the time spent in Mito_Cat_ and to a greater degree, Mito_Ox_ (Figure S3b), leading to a modest reduction in the duty cycle as well as a more significant reduction in the period of the oscillation (Figure 3f). These striking observations suggest that the PKm2-driven PEP cycle regulates the onset, and even more strongly, the termination of Ca^2+^ influx. Consistently, in PCK2-βKO islets where mitochondrial PEP production is inhibited, both the period and amplitude of glucose-stimulated Ca^2+^ oscillations were increased relative to controls islets (Figure 3h). Although the duty cycle also increased, it was only by a small margin. The period lengthening occurred from an increased duration of Mito_Cat_, and especially Mito_Ox_ (Figure S3c). Taken together, these data indicate that PKm2 controls both an “on-switch” and an “off-switch” for Ca^2+^ oscillations, both dependent on the mitochondrial production of PEP.

While above experiments examined conditions at a fixed elevated glucose concentration (10 mM), we also investigated Ca^2+^ dynamics following the transition from low to high glucose where first-phase insulin secretion is observed. Preincubation of control islets with PKa reduced the time to depolarization as well as the duration of the first-phase Ca^2+^ influx (Figure 3g). Conversely, depolarization was delayed in PCK2-βKO islets (Figure 3i). In this case, the duration of the first-phase Ca^2+^ pulse was not calculated since nearly 60% of PCK2-βKO islets failed to exit the first phase plateau in order to begin oscillations, as compared with only 27% of control islets (Figure 3i). In other words, while the PCK2 knockout had a weaker first phase Ca^2+^ rise, it had a much longer plateau that failed to turn off effectively. Hence, PKm2 activation and PCK2 serve as on-switches for promoting glucose-stimulated Ca^2+^ influx during the triggering phase (Mito_Cat_), and with a quantitatively larger effect, off-switches during the secretory phase (Mito_Ox_).

### Mitochondrial PEP carboxykinase (PCK2) is essential for amino acids to promote a rise in cytosolic ATP/ADP

Amino acids (AA) are obligate mitochondrial fuels that simultaneously feed oxidative and anaplerotic pathways. AA can be used as a tool for separating mechanistic components of the secretion mechanism because at low glucose they can, independently of glycolysis, raise ATP/ADP_c_ and elicit K_ATP_ channel closure, Ca^2+^ influx, and insulin release. In particular, glutamine and leucine generate PEP via glutamate dehydrogenase (GDH)-mediated anaplerosis that is followed by PCK2-mediated cataplerosis of PEP (Kibbey et al., 2014; Stark et al., 2009). We first examined whether restriction of mitochondrial PEP production in PCK2-βKO islets impacts the cytosolic ATP/ADP_c_ ratio, measured with β-cell specific expression of Perceval-HR biosensors. To limit glycolytic PEP, islets were incubated at 2.5 mM glucose. The islets were then stimulated with a mixture of AA including leucine and glutamine to allosterically activate and fuel GDH, respectively. Consistent with defective PEP cataplerosis, the ATP/ADP_c_ response of PCK2-βKO islets was only 37% of control islets in response to AA (Figure 4a). In this setting of PCK2 depletion, pharmacologic PK activation did not recover any of the AA-induced ATP/ADP_c_ response due to the absence of either a glycolytic or mitochondrial PEP source (Figure 4b). As expected, deletion of either PKm1 or PKm2 had only modest effects on the β-cell ATP/ADP_c_ response to AA (Figure 4c,e). However, PKa completely recovered the AA-induced rise in ATP/ADP_c_ in PKm1-βKO islets, in which the allosteric PKm2 isoform remains (Figure 4d), but not in PKm2-βKO islets (Figure 4f). Thus, while neither PK isoform is required for AA stimulation to increase ATP/ADP_c_, PCK2 displays high control strength over ATP/ADP_c_ in response to AA, especially considering that the PCK2 protein was only reduced by about two-thirds (Figure 1d).

**Figure 4.**
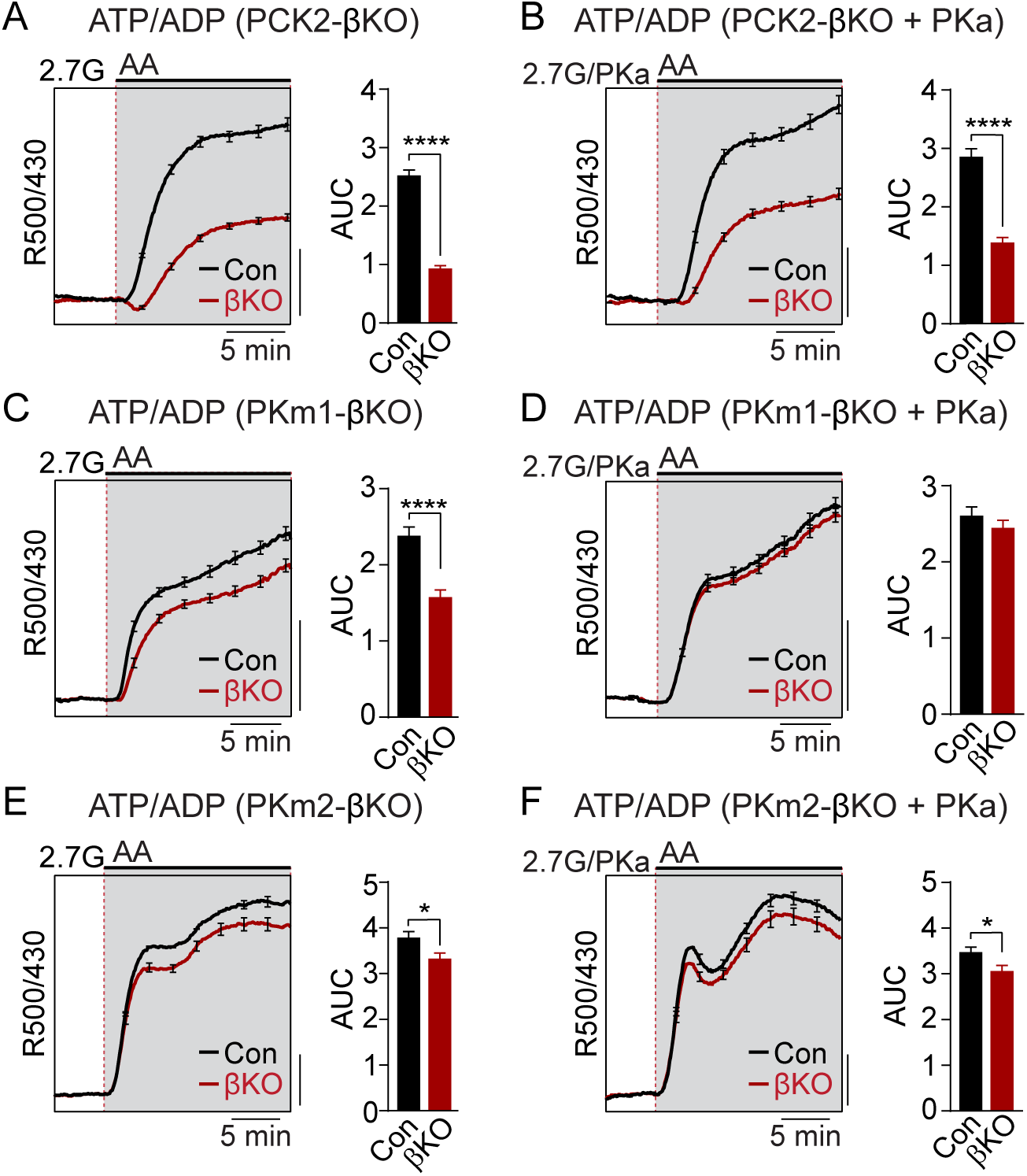
Restriction of the glycolytic PEP supply reveals the importance of PCK2 for cytosolic ATP/ADP. Average β-cell ATP/ADP in islets from PCK2-βKO (A-B), PKm1-βKO (B-E), and PKm2-βKO (E-F) mice in response to mixed amino acids (AA) provided at three times their physiological concentrations (1x = Q, 0.6 mM; L, 0.5 mM; R, 0.2 mM; A, 2.1 mM) in the presence of 2.7 mM glucose (2.7G) to remove the enolase contribution to cytosolic PEP. PKa (10 μM TEPP-46) was present in B, D, and F. ATP/ADP is quantified as area under the curve (AUC) from PCK2-βKO (vehicle, *n* = 73 islets from 3 mice; PKa, *n* = 71 islets from 3 mice) and littermate controls (vehicle, *n* = 90 islets from 3 mice; PKa, *n* = 77 islets from 3 mice); PKm1-βKO (vehicle, *n* = 66 islets from 3 mice; PKa, *n* = 72 islets from 3 mice) and littermate controls (vehicle, *n* = 69 islets from 6 mice; PKa, *n* = 69 islets from 3 mice); and PKm2-βKO (vehicle, *n* = 99 islets from 3 mice; PKa, *n* = 89 islets from 3 mice) and littermate controls (vehicle, *n* = 100 islets from 6 mice; PKa, *n* = 85 islets from 3 mice). Scale bars: 0.1 Perceval-HR excitation ratio (R500/430). Data are shown as mean ± SEM. \**P* < 0.05, **** *P* < 0.0001 by t-test.

### Mitochondrial fuels that stimulate a bulk rise in ATP/ADP_c_ fail to close K_ATP_ in the absence of PK

Mitochondria are located throughout the β-cell, including near the plasma membrane, where submembrane ATP microdomains have been observed (Griesche et al., 2019; Kennedy et al., 1999). Since some PK is located on the plasma membrane, we wondered whether during AA stimulation mitochondria can provide PEP to facilitate PK-dependent K_ATP_ closure, or alternatively, whether mitochondria can serve as a direct source of ATP for K_ATP_ channel closure. To determine whether mitochondrial PEP impacts the K_ATP_ channel microcompartment, we monitored K_ATP_ channel currents in *intact* β-cells in the cell-attached configuration in response to bath-applied AA at 2.5 mM glucose (Figure 5a). Mixed AA with or without PKa reduced K_ATP_ channel power (reflecting the total number of transported K^+^ ions) by ∼75% in control β-cells (Figure 5b). However, no K_ATP_ closure was observed in the absence of PCK2, even with PK activator present (Figure 5c). These findings indicate that mitochondrially-derived PEP can signal to the K_ATP_ channel microcompartment, and is essential for K_ATP_ closure in response to AA.

**Figure 5.**
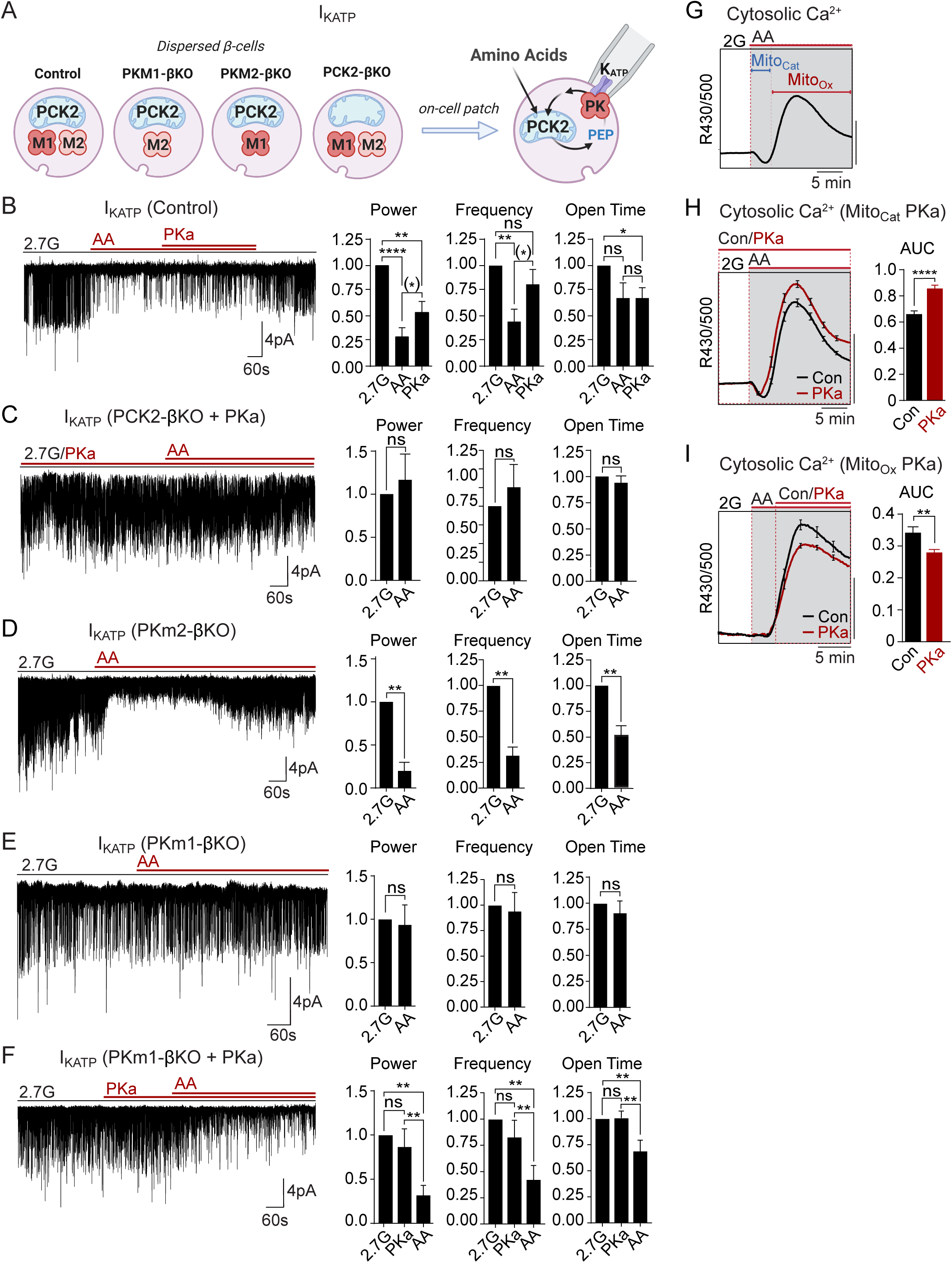
Mitochondrial PEP signals to PK within the plasma membrane K_ATP_ channel microcompartment in intact β-cells. (A) Diagram of on-cell patch clamp method in intact β-cells with bath application of amino acids.(B-F) Representative example traces and quantification of K_ATP_ channel closure in terms of normalized power, frequency, and open time for β-cells from control (*n* = 10 recordings from 3 mice) (A), PKm1-βKO (*n* = 7 recordings from 3 mice) (B-C), PKm2-βKO (*n* = 7 recordings from 3 mice) (E), and PCK2-βKO mice (*n* = 10 recordings from 3 mice) (F) in response to mixed amino acids (AA) and 2.7 glucose (2.7G) as in Figure 4. (G-I) AA-stimulated Ca^2+^ responses in control islets illustrating the Mito_Cat_ and Mito_Ox_ phases (A). The average Ca^2+^ response to PKa application during Mito_Cat_ (vehicle, *n* =575 islets from 8 mice; PKa, *n* = 575 islets from 8 mice) (H) and Mito_Ox_ (vehicle, *n* = 91 islets from 4 mice; PKa, *n* = 108 islets from 4 mice) (I) is quantified as AUC. Scale bars: 0.025 FuraRed excitation ratio (R430/500). Data are shown as mean ± SEM. \**P* < 0.05, \*\**P* < 0.01, \*\*\**P* < 0.001, **** *P* < 0.0001 by paired one-way ANOVA or paired t-test as appropriate. Following the removal of one outlier by ROUT (Q=10%), ^(*)^*P* < 0.05 by paired t-test in (B).

Unlike PCK2-deficient β-cells, PKm1-deficient β-cells stimulated with AA were capable of increasing ATP/ADP_c_ to a similar level as controls (Figure 4). This model provides a unique opportunity to directly test the canonical model of fuel induced insulin secretion, where a rise in ATP/ADP in the bulk cytosol is thought to be sufficient to close K_ATP_ channels. While K_ATP_ channels were efficiently closed by AA in control and PKm2-deficient β-cells (Figure 5b and 5d), K_ATP_ channels failed to close in β-cells lacking PKm1 (Figure 5e). As in excised patches (Figure 2c-d), pharmacologic activation of PKm2 was sufficient to rescue K_ATP_ closure in PKm1-deficient β-cells (Figure 5f). These remarkable findings demonstrate that PK activity is essential for K_ATP_ channel closure in response to AA, and argue strongly against the canonical model in which mitochondrially-derived ATP raises ATP/ADP_c_ to close K_ATP_ channels.

### Mitochondrially-derived PEP drives K_ATP_ closure and Ca^2+^ influx during Mito_Cat_, and accelerates K_ATP_ reopening and Ca^2+^ extrusion during Mito_Ox_

As for glucose, AA-stimulated Ca^2+^ influx follows a distinct triggering and secretory phase, representing Mito_Cat_ and Mito_Ox_, respectively (Figure 5g). Following K_ATP_ channel closure with AA, the surprising ability of PKa to partially reopen K_ATP_ channels (Figure 5b) suggests that PKm2 activation might also serve as an “off-switch” that ultimately increases the channel opening by activating the PEP cycle. This entirely novel concept is consistent with the with the ability of PKa to both hasten the onset *and* shorten the duration of Ca^2+^ pulses (Figure 3f,g). It is also consistent with the observation that mitochondrially-derived PEP is necessary to switch off glucose-dependent Ca^2+^ influx, as shown in PCK2-βKO islet experiments (Figure 3h,i). In other words, the PEP cycle would have a dual function in the β-cell: during Mito_Cat_, the PEP cycle facilitates K_ATP_ closure and Ca^2+^ influx; during Mito_Ox_, the PEP cycle may facilitate K_ATP_ channel reopening, Ca^2+^ extrusion and turn off insulin secretion. To test this concept further, we examined the effect of PKa on AA-stimulated Ca^2+^ influx. When applied *before* AA stimulation, during Mito_Cat_, PKa increased Ca^2+^ influx (Figure 5h). By contrast, PKa application *after* the initial Ca^2+^ rise, during Mito_Ox_, reduced cytosolic Ca^2+^ (Figure 5i). These data confirm two temporally separated functions of PK in response to mitochondrial PEP – representing on and off-switches for β-cell Ca^2+^.

### PKm1 and PKm2 respond differentially to the glycolytic and mitochondrial sources of PEP

We next examined the functional consequence of β-cell PKm1, PKm2, and PCK2 deletion on AA-stimulated Ca^2+^ influx and insulin secretion at both low and high glucose (Figure 6). In PCK2-βKO islets at 2 mM glucose, the AA-induced Ca^2+^ response was reduced along with insulin secretion (Figure 6a,b). In the presence of 10 mM glucose, β-cell PCK2 deletion did not impact insulin secretion because of the restored glycolytic PEP supply (Figure 6c). At high glucose, PCK2-βKO islets fail to inactivate during Mito_Ox,_ as indicated by the sustained Ca^2+^ plateau (Figure 3h,i). Thus, when anaplerosis was maximally stimulated by 10 mM glucose and AA, insulin secretion was higher in PCK2-βKO islet than controls (Figure 6c).

**Figure 6.**
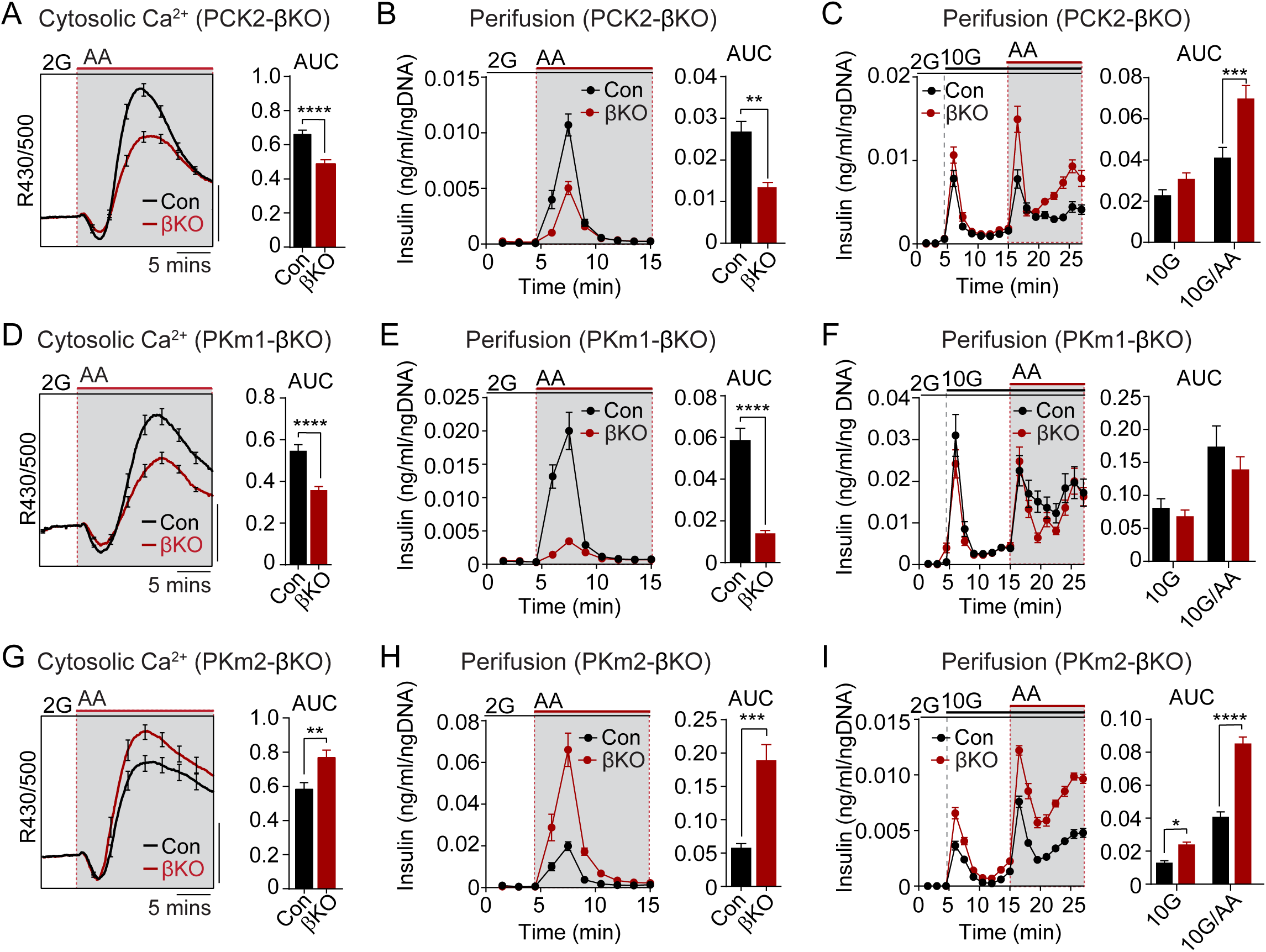
The PKm1/PKm2 ratio dictates the β-cell Ca_2+_ and secretory response to anaplerotic fuels. Average cytosolic Ca^2+^ and insulin secretory responses in controls and PCK2-βKO (A-C), PKm1-βKO (D-F) and PKm2-βKO islets (G-I) to the indicated concentrations of glucose (2G, 2 mM; 10G, 10 mM) and amino acids (AA; concentrations are listed in Figure 4). Data are quantified as AUC. Islet Ca^2+^ data reflect *n* = 60-101 islets from 3 mice per condition (scale bar = 0.025 FuraRed excitation ratio). Islet perifusion data reflect *n* = 100 islets per mouse and 6 mice per condition except PKm2-βKO at high glucose (5 mice). Data are shown as mean ± SEM. *p<0.05, **p<0.01, ***p<0.001, ****p<0.0001 by t-test.

Like the PCK2-βKO, the Ca^2+^ and secretory responses of PKm1-βKO islets were blunted in AA-stimulated islets at 2 mM glucose (Figure 6d,e). However, the insulin secretory response remained intact at 10 mM glucose in PKm1-βKO islets, whether or not AA were present (Figure 6f), due to the sufficiency of PKm2 in the presence of glycolytic FBP. These findings are entirely consistent with the differential response of PKm1-βKO islets to AA vs. glucose stimulation observed in the Ca^2+^ and K_ATP_ channel recordings above (Figures 3 and 5).

Similarly to PCK2, allosterically-activated PKm2 has dual functions during Mito_Cat_ and Mito_Ox_, as evidenced by the Ca^2+^ and K_ATP_ channel measurements shown in Figures 3f, 3h, 5b, 5h, and 5i. Both the AA-induced Ca^2+^ response and insulin secretion were greatly increased in PKm2-βKO islets compared to controls (Figure 6g,h). Glucose alone, but especially in combination with AA, stimulated enhanced secretion in the absence of PKm2 (Figure 6i). Thus, while either PKm1 or PKm2 is sufficient to initiate glucose-stimulated insulin secretion during Mito_Cat_, PKm2 is essential for mitochondrial PEP to switch the system off during Mito_Ox_. When PKm2 was deleted, and PKm1 increased (Figure 1c), the system is shifted towards greater nutrient-induced and PCK2-dependent insulin secretion with heightened sensitivity to anaplerotic fuels.

## DISCUSSION

These data provide genetic evidence that PEP has metabolic control over PK-dependent ATP/ADP_pm_ generation, K_ATP_ closure, Ca^2+^ signaling, and insulin secretion. β-cell PK isoform deletion experiments demonstrate that plasma membrane-associated PK is strictly required for K_ATP_ channel closure and provide rigorous genetic evidence for the Mito_Cat_-Mito_Ox_ model of oscillatory metabolism and insulin secretion. Our results indicate that PK is controlled by two different sources of PEP – glycolytic and mitochondrial (Figure 1a). Glucose signals to PK via both glycolytic and mitochondrially-derived PEP, whereas amino acids signal exclusively through mitochondrially-derived PEP. Since amino acids do not generate FBP, which is needed to allosterically activate PKm2, PKm1 is necessary to raise ATP/ADP_pm_, close K_ATP_ channels, and stimulate insulin secretion in response to amino acids. Furthermore, our work supports the concept that the canonical model of fuel-induced insulin secretion, whereby mitochondrially-derived ATP produced via the electron transport chain raises ATP/ADP_c_ to close K_ATP_ channels, is possibly wrong. The evidence for this new view of β-cell metabolic signaling is that amino acids that efficiently raise ATP/ADP_c_ do not close K_ATP_ channels in the absence of plasma membrane PK activity. Finally, the data indicate that PK and the PEP cycle have a dual role in the control of insulin secretion – they act as on-signals during the triggering phase, Mito_Cat_, and as off-signals during the active secretory phase, Mito_Ox_. We discuss each of these findings in the sections below.

An amino acid-stimulated rise in ATP/ADP_c_ was shown to be insufficient to close K_ATP_ channels, ruling out a key aspect of the “canonical model” in which accelerated mitochondrial metabolism raises ATP/ADP_c_ to close K_ATP_ channels (Campbell and Newgard, 2021; Prentki et al., 2013; Thompson and Satin, 2021). In PKm1-deficient β-cells, amino acids increased ATP/ADP_c_ similarly to control β-cells, but were unable to close K_ATP_ channels. This defect was rescued by pharmacological re-activation of PKm2, present on the plasma membrane of PKm1-deficient β-cells, which acutely restored K_ATP_ closure in response to amino acids. Thus, PK is essential for K_ATP_ closure in intact β-cells. Taken together with the sufficiency of plasma membrane PK for K_ATP_ closure in excised patch experiments, our data support the Mito_Cat_-Mito_Ox_ model in which plasma-membrane compartmentalized PK increases the ATP/ADP ratio to *initiate* insulin secretion, while oxidative phosphorylation plays the dominant role in providing ATP to sustain insulin secretion *after* membrane depolarization (Lewandowski et al., 2020).

In the mouse β-cell, we demonstrate that most of the PK activity is the constitutively-active PKm1 isoform, with a minor contribution from PKm2 and no detectable PKL activity. In the rat, MacDonald and Chang (MacDonald and Chang, 1985) reported PK activity to be primarily PKm2, which may be a species difference. However, in this study FBP was shown to have a minor effect on PK activity, which casts doubts about the conclusion of this work. Our prior studies of PK-dependent K_ATP_ closure in human β-cell plasma membranes were conducted in the absence of FBP, which is consistent with the presence of PKm1 in the microdomain but not ruling out PKm2 (Lewandowski et al., 2020). Regardless, PKm2 activators amplify nutrient-stimulated insulin secretion in mouse, rat, and human islets *in vitro*, and in rats *in vivo*, indicating that PKm2 is functionally recruitable in all three species (Abulizi et al., 2020; Lewandowski et al., 2020).

Whole-body PCK2^-/-^ mice display glucose intolerance and reduced insulin secretion in response to glucose and amino acids (Abulizi et al., 2020). These effects may arise independently of the β-cell, since here we show that PCK2-βKO mice of similar age are glucose tolerant, and glucose-stimulated insulin secretion from isolated islets is actually increased when amino acids are present. Nevertheless, Ca^2+^ influx induced by amino acid stimulation at low glucose is blocked in both models. We further demonstrate that, in response to amino acids, β-cell PCK2 is rate controlling for ATP/ADP_c_, K_ATP_ closure, Ca^2+^ influx and insulin secretion. However, in the presence of elevated glucose, amino acid stimulation in fact increased insulin secretion. This may be explained by the hypothesized dual function of the PEP cycle playing a role in both the on- and off-phases of GSIS. β-cell deletion of PCK2 had a strong effect on glucose-stimulated Ca^2+^ oscillations, delaying *both* the onset of Ca^2+^ influx and preventing the ability of β-cells to efficiently repolarize. The failure of this off-switch provides an explanation for the hypersecretion in β-cells lacking PCK2 when anaplerosis is fully primed with the combined presence of glucose and amino acids.

Insulin release is frequently described in terms of triggering and amplifying on-signals termed “metabolic coupling factors” (MCFs). As defined by Prentki, “regulatory MCFs” are nutrient-dependent signals that facilitate the switch between metabolic networks (e.g. malonyl-CoA switches β-cells from fatty acid to glucose oxidation), while “effectory MCFs” (e.g., Ca^2+^, ATP/ADP, monoacylglycerol, and reactive oxygen species), are transient, necessary, and sufficient on-signals that dose-dependently stimulate insulin secretion (Prentki et al., 2013). Is PEP a regulatory or effectory signal for insulin secretion – or both? PEP has some properties of a regulatory signal, since its metabolism by PK generates a bioenergetic feed forward that progressively deprives mitochondria of ADP, shutting down oxidative phosphorylation in favor of activating both pyruvate carboxylase and the PEP cycle (Lewandowski et al., 2020; Sugden and Ashcroft, 1977). Through a positive Hill coefficient, the increasing concentration of PEP progressively activates PKm2 to increase ATP/ADP (Lewandowski et al., 2020; Merrins et al., 2013), further reinforcing the PEP cycle. ATP/ADP_pm_ is clearly an effectory signal in that it is sufficient to cause depolarization (Ashcroft et al., 1984; Cook and Hales, 1984; Rorsman and Trube, 1985) while at the same time priming granule exocytosis (Eliasson et al., 1997; Pizarro-Delgado et al., 2016; Takahashi et al., 1999). Arguably, PEP also acts as an effectory MCF, since its presence in the K_ATP_ microenvironment in the excised patches can override channel opening by continuous ADP to close K_ATP_ channels. This property requires either PKm1 or allosterically-activated PKm2.

Effectory MCFs must be counterbalanced by a strong off-switch that ensures the signal is transient. That is, β-cells can fail if K_ATP_-dependent Ca^2+^ influx is activated without a coordinated homeostatic mechanism to turn it off (Remedi and Nichols, 2009). While PKm2 activation during Mito_Cat_ was found to facilitate K_ATP_ closure and increase Ca^2+^, we also found that PKm2 activation during Mito_Ox_ accelerated K_ATP_ channel reopening and lowered Ca^2+^. Like the on-switch, this off-switch may also involve mitochondrial PEP, since β-cell PCK2 deletion stalls the β-cell in the Ca^2+^-activated state. In addition to these temporally compartmentalized effects, spatial considerations, such as the stimulus-dependent movement of the mitochondria away from the plasma membrane (Griesche et al., 2019), may also be important for Ca^2+^ oscillations. Further studies are needed to elucidate precisely how the PK and the PEP cycle may contribute to turning off Ca^2+^ influx and insulin secretion during the Mito_Ox_ phase. Finally, it remains to be determined whether β-cell PK and the PEP cycle are altered in the states of obesity and diabetes, and whether they contribute to hyperinsulinemia and/or β-cell failure.

## ACKNOWLEDGEMENTS

We would like to thank Matthew Vander Heiden (MIT) for providing PKm1^f/f^ mice, and Mark Huising (UC-Davis) and Barak Blum (UW-Madison) for providing Ucn3-Cre mice. We would also like to acknowledge Kathy Krentz and Dustin Rubenstein at the UW-Madison Genome Editing and Animal Model Core for their assistance generating PCK2^f/f^ mice, and Jiwon Seo and Jody Peter at the UW-Madison Biomedical Research Model Services Breeding Core for their assistance with animal husbandry and genotyping. Graphics were created using BioRender.com. The Merrins laboratory gratefully acknowledges support from the NIH/NIDDK (R01DK113103 and R01DK113103) and the Department of Veterans Affairs (I01B005113). HRF received a postdoctoral fellowships from HRSA (T32HP10010) and the NIH/NIA (T32AG000213), SLL received a predoctoral fellowship from the NIH/NIDDK (T32DK007665), HRV received a postdoctoral fellowship from the American Diabetes Association (1-17-PDF-155), and we acknowledge Dudley Lamming for contributing support for EP from R01AG062328. The Kibbey laboratory gratefully acknowledges support from the NIH/NIDDK (R01DK127637). This work utilized facilities and resources from the William S. Middleton Memorial Veterans Hospital and does not represent the views of the Department of Veterans Affairs or the United States Government.

## AUTHOR CONTRIBUTIONS

MJM conceived the study and wrote the paper with HRF, MP and RGK. HRF performed the main body of the experiments with assistance from TH, EP, SMS, SLL, HRV, SMD, and RLC. MJM and RGK provided resources. All authors interpreted the data and edited the manuscript.

## MATERIALS AND METHODS

### Mice Creation

β-cell specific PKm1 and PKm2 knockout mice were generated by breeding *Ins1-Cre* (Thorens et al., 2015) mice (B6(Cg)-*Ins1*^*tm1*.*1(cre)Thor*^*/J*, Jackson Laboratory #026801) with *Pkm1*^*f/f*^ mice (Davidson et al., 2021; Li et al., 2021, p. 1) provided by Matthew Vander Heiden (MIT) or *Pkm2*^*f/f*^ mice (Israelsen et al., 2013) (B6;129S-*Pkm*^*tm1*.*1Mgvh*^/J, Jackson Laboratory #024408) after 10 generations of backcrossing to C57BL6/J mice (Jackson Laboratory). *Pck2*^*f/f*^ mice were generated *de novo* by the University of Wisconsin-Madison Genome Editing and Animal Model core. Backcrossed F1s were sequence confirmed by Illumina targeted deep sequencing to confirm the LoxP insertions around exon 5 (ENSMUSE00000399990; Figure S1), and the intervening region was sequence confirmed with Sanger sequencing. *Pck2*^*f/f*^ mice were crossed with *Ucn3-Cre* mice (van der Meulen et al., 2017) provided by Barak Blum (University of Wisconsin-Madison) with permission from Mark O. Huising (University of California-Davis). All mice were genotyped by Transnetyx.

### Mouse Islet Preparations

Male mice were housed 2-5 per cage at 21-23°C, fed with a chow diet and water provided *ad libitum*, and maintained on a 12 h light/dark cycle. Mice 12-20 weeks of age were sacrificed via CO_2_ asphyxiation followed by cervical dislocation, and islets were isolated using methods previously described (Gregg et al., 2016).

### Cloning and Adenoviral Delivery of Biosensors

Generation of adenovirus carrying genetically-encoded ATP/ADP biosensors (Perceval-HR) under control of the insulin promoter was described previously (Merrins et al., 2016). High-titer adenovirus was added to islets immediately after islet isolation and incubated for 2 h at 37°C then moved to fresh media. Imaging was performed 3 days post isolation.

### Western blots

Islets were lysed using 0.1% Triton-X100 in phosphate buffered saline (1 μl/islet). Islets in lysis buffer were incubated at room temperature for 15 minutes, vortexed for 30 seconds, frozen/thawed, and vortexed for 30 seconds again. Cell lysate was spun down at max speed in a table top centrifuge and 16-20 μl supernatant was added to each well of a 12% SDS-PAGE gel. The gel was run at 110V for ∼2 hours then transferred to a PVDF membrane at 100V for 1 hour at 4°C. Membranes were blocked in 4% BSA in Tris-buffered saline containing 0.1% Tween-20 detergent (TBST) for 30 minutes then incubated overnight with PKm1 (Cell Signaling #4053; 1:1000), PKm2 (Cell Signaling #7067; 1:1000), PCK2 primary antibodies (Cell Signaling #6924; 1:1000), or loading controls beta-actin (Cell Signaling #3700; 1:1000) or HSP90 (Cell Signaling #4877; 1:1000). Blots were washed for 15 min in TBST four times then incubated for 2 hours at room temperature with an anti-rabbit IgG HRP-linked secondary antibody (Cell Signaling #7074; 1:20,000). Blots were imaged and quantified using ImageQuant (Amersham).

### Timelapse Imaging

Islets were preincubated in 2.5 μM FuraRed (Molecular Probes F3020) in RPMI 1640 for 45 minutes at 37°C prior to being placed in an glass-bottomed imaging chamber (Warner Instruments) on a Nikon Eclipse Ti-E inverted microscope with a Super Flour 10x/0.5N.A. objective (Nikon Instruments). The chamber was perfused with standard imaging solution (in mM: 135 NaCl, 4.8 KCl, 5 CaCl_2_, 20 HEPES, 1.2 MgCl_2_) for low glucose/amino acid experiments, or HBSS (in mM 137 NaCl, 5.6 KCl, 1.2 MgCl_2_, 0.5 NaH_2_PO_4_, 4.2 NaHCO_3_, 10 HEPES, 2.6 CaCl_2_) for high glucose experiments containing glucose and amino acid concentrations indicated in figure legends. Temperature was maintained at 33°C with inline solution and chamber heaters (Warner Instruments), and flow rate was set to 0.25 ml/min. Excitation was provided by a SOLA SE II 365 (Lumencor). A Hamamatsu ORCA-Flash4.0 V2 Digital CMOS camera was used to collect fluorescence emission at 0.125-0.2 Hz. Nikon NIS-Elements was used to designate regions of interest. Excitation (x) and emission (m) filters (ET type; Chroma Technology) were used with an FF444/521/608-Di01 dichroic (Semrock) as follows: FuraRed, 430/20x and 500/20x, 630/70m (R430/500 was reported) and Perceval-HR, 430/20x and 500/20x, 535/35m (R500/430 was reported). A custom MATLAB script was used to quantify cytosolic Ca^2+^ oscillation parameters (script available at https://github.com/hrfoster/Merrins-Lab-Matlab-Scripts). In order to differentiate between islets of control and knockout mice, islets of one genotype were barcoded by pre-incubation with 2 μM DiR (Fisher Scientific D12731) or 2 μM DiD (Fisher Scientific D7757) in islet media for 10 minutes at 37°C. Islet barcodes were then imaged using a Cy7 (for DiR) or Cy5 filter cube (for DiD) from Chroma.

### Electrophysiology

Single channel patch clamp experiments and data analysis were performed as previously described (Lewandowski et al., 2020). Briefly, mouse and human cells were acutely isolated by dispersing of islets with Accutase (Fisher Scientific). Cells were plated on sterilized uncoated glass shards and kept at 37°C with 95% O_2_-5% CO_2_. For both on-cell and inside-out recordings, gigaseals were obtained in extracellular bath solution (in mM): 140 NaCl, 5 KCl, 1.2 MgCl_2_, 2.5 CaCl_2_, 0.5 Glucose, 10 HEPES, pH 7.4, adjusted with NaOH and clamped at −50mV. For inside-out configuration, after pipette excision, the bath solution was replaced with equilibrium solutions with K^+^ as the charge carrier (in mM): 130 KCl, 2 CaCl_2_, 10 EGTA, 0.9 [Mg^2+^]_free_, 10 sucrose, 20 HEPES, pH 7.2 with KOH. The [Mg^2+^]_free_, calculated using WEBMAXC standard, was held constant in the presence of nucleotides. The pipette solution, used for both on-cell and inside-out configurations, contained (in mM): 10 sucrose, 130 KCl, 2 CaCl_2_, 10 EGTA, 20 HEPES, pH 7.2, adjusted with KOH. Recording electrodes made of microfilament borosilicate glass (Harvard Apparatus, Holliston, MA #64-0792) were used to pull patch pipettes (3 MΩ) on a horizontal Flaming/Brown micropipette puller (P-1000, Sutter Instruments) and polished by microforge (Narishige MF-830) to a final tip resistance of 5-10 MΩ. On-cell recording started after formation of a stable gigaseal (>2.5 GΩ) and inside-out recording started after withdrawal of the pipette and establishment of the excised inside-out configuration. A HEKA Instruments EPC10 patch-clamp amplifier was used for registration of current. Data was Bessel filtered online at 1 kHz and single channel currents were analyzed offline using ClampFit analysis module of pCLAMP 10 software (Molecular Devices).

### Islet Perifusion Assays

Islets from 6 mice of each genotype were pooled and then divided in equal numbers and placed into a 12-channel perifusion system (BioRep Peri-4.2; 75-100 islets/chamber) containing KRPH buffer (in mM: 140 NaCl, 4.7 KCl, 1.5 CaCl_2_, 1 NaH_2_PO_4_, 1 MgSO_4_, 2 NaHCO_3_, 5 HEPES, 2 glucose, 0.1% fatty acid-free BSA, pH 7.4) with 100 μl Bio-Gel P-4 Media (Bio-Rad). Islets were equilibrated at 2 mM glucose for 36-48 minutes prior to perfusion with amino acids or 10 mM glucose at 37ºC. Insulin secretion was assayed using Promega Lumit™ Insulin Immunoassay (CS3037A01) and measured using a TECAN Spark plate reader. Quant-IT PicoGreen dsDNA Assay Kit (Thermo Scientific P7589) was used to determine DNA content after lysis using 0.1% Triton X-100.

### PK activity

The enzymatic activity of pyruvate kinase was measured using EC 2.7.1.40 from Sigma (Bergmeyer, H.U. et al., n.d.) as described previously (Lewandowski et al., 2020). FBP (Sigma Aldrich #F6803) and TEPP-46 (PKa; Fisher Scientific #50-548-70001) were used at a concentration of 80 μM and 10 μM, respectively. Experiments were performed at 37ºC.

### Quantification and Statistical Analysis

Figure legends describe the statistical details of each experiments. Data are expressed as mean ± SEM. Statistical significance was determined by two-way or one-way ANOVA with Sidak multiple-comparisons test post hoc or Student’s t-test as indicated in the figure legends. Data were continuous and normally distributed so were analyzed with parametric tests. Differences were considered to be significant at P < 0.05. Calculations were performed using GraphPad Prism.

### Study approval

Animal experiments were conducted under the supervision of the IACUC of the William S. Middleton Memorial Veterans Hospital (Protocol: MJM0001).

