## Supplemental Information for "The isoforms of pyruvate kinase act as nutrient sensors for the β-cell K_ATP_ channel"

Supplemental Figure S1

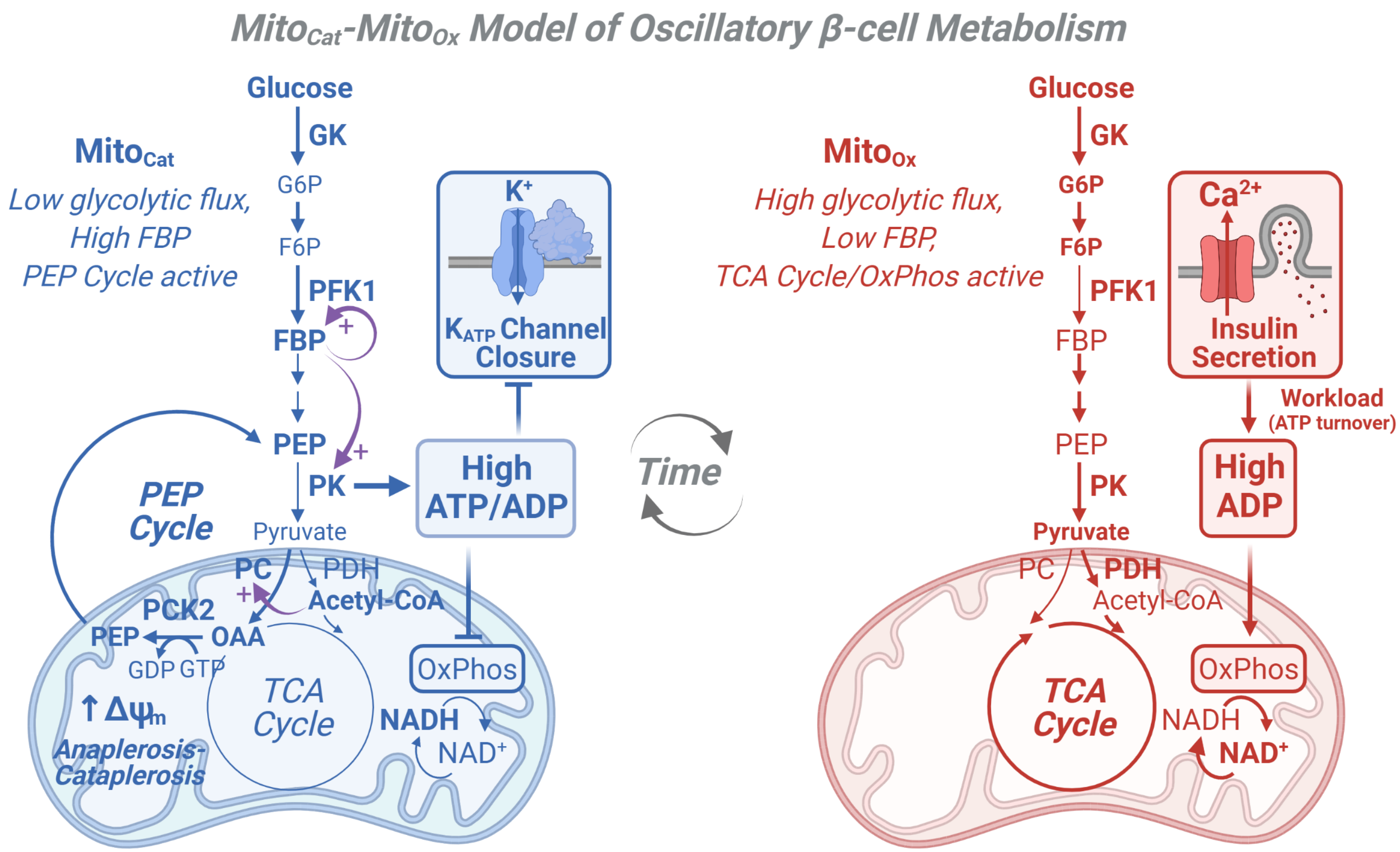

**Figure S1. Cartoon depicting the $Mito_{Cat}$-$Mito_{Ox}$ model of oscillatory β-cell metabolism.**

$Mito_{Cat}$-$Mito_{Ox}$ model of oscillatory β-cell metabolism with two states (triggering vs. secretory) separated by membrane depolarization. $Mito_{Cat}$ is interchangeable with $Mito_{Synth}$ in Lewandowski et al. (2020)[1] and is a more precise term reflecting the matched processes of anaplerosis and cataplerosis that are central to glucose signaling. $Mito_{Cat}$: Before depolarization, PK lowers ADP, reducing oxidative phosphorylation (OxPhos) and the TCA cycle, which raises acetyl-CoA that allosterically activates pyruvate carboxylase and PEP cataplerosis through PCK2. The return of PEP to the cytosol completes the PEP cycle and augments PK. $Mito_{Ox}$: After PK raises ATP/ADP sufficiently to close $K_{ATP}$ channels, workload in the form of ATP hydrolysis restores ADP (e.g. by exocytosis and pumps), increasing OxPhos, TCA cycle, and glycolytic flux. Note that recruitable PK isoforms (M2 and L) are active before depolarization, when glycolytic flux is low and the fructose 1,6-bisphosphate (FBP) concentration is high.

### Supplemental Figure S2

AAATCTTACCCATCCACGGAGAAATTGTGGATGTAGAACGATTATTCC**ATAACTTCGTATAATGTATGCTATACGA**
**AGTTAT**CCGAGGCCCAGTGACGCATTCTGGACTGGAGTAACACCATTTGGAGAACAGAAGTCAGTCGACAGGTCT
GACAAAGCACGAGGGCTGATGGGTACTGAAGAAGATGGTTTAGCTACCCCTGGGCATTCTATTGCTTCCAAGTGT
CTCTTGGGTCAATCCTTGATCCCTCCAGTCCCTGACACCCTGGTTTCTGATGCCAGTGGCAGCAGCACTATGACCCC
ATTGTCTCCAGGGGATCCTGTGGGCCAGTGGCCGTGCAATCCAGAAAAAACCCTGATTGGCCACGTGCCAGACCA
GCGGGAGATCGTCTCCTTCGGCAGCGGCTATGGTGGTAACTCCTTGCTGGGCAAGAAGTGCTTTGCCCTGCGCAT
CGCCTCTCGCCTGGCCAGGGATGAGGGCTGGCTGGCAGAGCACATGCTGGTGAGGGCCTGGTGAGGAACTGAGC
AGCTGTGGTAGGGGAGTGGGTGGGGAAGCCTTGGCAATCTGCCTCAGCTTTGCCTCCTTCCTGCCAGGTGCCAGG
AGGATGAGCCTGAAACCTGAAGTTTTAGC**ATAACTTCGTATAATGTATGCTATACGAAGTTAT**CCCAGGCTGCTG
AGGTGCTACCCCAGTCTCTAGCGGGGCAGGACACCTG

**Figure S2. Sequence verification of PCK2$^{f/f}$ mice related to Fig. 1.** Genomic DNA sequence of *pck2* around exon 5 (ENSMUSE00000399990; in red), showing LoxP sites (bold) as confirmed by Illumina targeted deep sequencing.

### Supplemental Figure S3

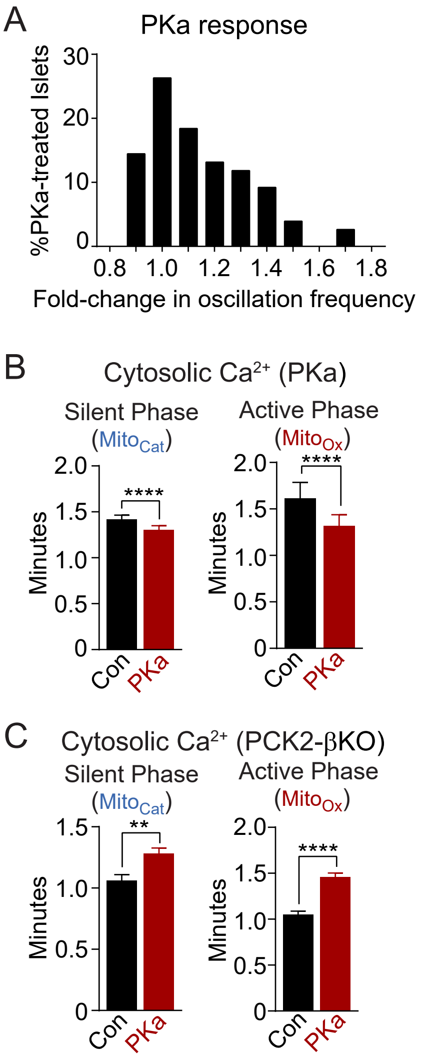

**Figure S3. Supplemental quantification of steady-state $Ca^{2+}$ dynamics related to Fig. 3.**

(A) The steady-state $Ca^{2+}$ response of mouse islets to PK activator (PKa, 10 μM TEPP-46) shows a primarily one-tailed distribution towards reduced period. 10 mM glucose. *n* = 76 islets from 3 mice.

(B) Quantification of the average duration of the silent phase ($Mito_{Cat}$) and active phase ($Mito_{Ox}$) of steady-state islet $Ca^{2+}$ oscillations from control mice with and without PKa in the presence of 10 mM glucose and 1 mM leucine (PKa, 10 µM TEPP-46). Control -PKa *n* = 88 from 3 mice, control +PKa *n* = 92 from 3 mice.

(C) Quantification of the average duration of the silent phase ($Mito_{Cat}$) and active phase ($Mito_{Ox}$) of steady-state islet $Ca^{2+}$ oscillations from PCK2-βKO mice and littermate controls in the presence of 10 mM glucose and 1 mM leucine. Control, *n* = 74 islets from 3 mice; PCK2-βKO, *n* = 77 islets from 3 mice. Data are shown as mean ± SEM. **p<0.01, ****p<0.0001 by unpaired t-test (S3D).
